# Engineering sorghum for higher 4-hydroxybenzoic acid content

**DOI:** 10.1101/2021.07.13.452095

**Authors:** Chien-Yuan Lin, Yang Tian, Kimberly Nelson-Vasilchik, Joel Hague, Ramu Kakumanu, Mi Yeon Lee, Jessica Trinh, Trent R. Northen, Edward E. K. Baidoo, Albert P. Kausch, Henrik V. Scheller, Aymerick Eudes

## Abstract

Lignocellulosic biomass represents a renewable source of sugars for the manufacturing of bioproducts such as biofuels. The high cost associated with deconstruction of plant biomass to simple sugars remains one of the challenges preventing the deployment of economically sustainable advanced bioproducts. The accumulation *in-planta* of value-added coproducts such as platform chemicals can improve the economics of biofuels. Among other crops, sorghum is an ideal bioenergy feedstock due to its low input requirements, efficient nitrogen recycling, and high water use efficiency and biomass yields. In this work, we engineered sorghum to overproduce the valuable chemical 4-hydroxybenzoic acid (4-HBA) by co-expressing plastid-targeted versions of *Escherichia coli* chorismate pyruvate-lyase (UbiC) and feedback-resistant 3-deoxy-D-arabino-heptulonate-7-phosphate synthase (AroG*). Two independent lines containing the dual *aroG\**-*ubiC* construct were selected for characterization in the T2 generation. Using liquid chromatography-mass spectrometry, analysis of methanolic extracts obtained from biomass samples revealed an accumulation of 4-HBA in the two transgenic lines (corresponding to 1.56% and 1.72% dry weight, respectively), with 4-HBA glucose conjugates representing major forms. Measurements of biomass composition and several agronomic traits showed no significant difference between transgenic and wild-type control plants grown under controlled environment. This work demonstrates the transferability of the *ubiC* engineering approach to sorghum; generated lines will be useful to assess the agronomic performance of modified 4-HBA-rich sorghum under natural conditions.

## Results and Discussion

Lignocellulosic biomass constitutes a renewable source of sugars for the manufacturing of bioproducts such as biofuels. The high cost associated with deconstruction of plant biomass to simple sugars remains one of the challenges preventing the deployment of economically sustainable advanced bioproducts. One proposed solution to improve the economics of biofuels is the *in-planta* accumulation of value-added coproducts (Yang et al., 2020). In this scenario, engineered bioenergy crops not only provide carbohydrates for conversion into bioproducts but also produce valuable compounds such as polymers, platform chemicals, pharmaceuticals, flavors and fragrances (Lin and Eudes, 2020).

Sorghum is an ideal bioenergy feedstock due to its low input requirements, efficient nitrogen recycling, and high water use efficiency and biomass yields (Mullet et al., 2014). Although several studies reported on the genetic modification of sorghum to reduce its recalcitrance to deconstruction, there are only a few examples of metabolic engineering for accumulation of bioproducts in sorghum biomass (Vanhercke et al., 2018). In this work, we tested the feasibility of engineering sorghum to overproduce 4-hydroxybenzoic acid (4-HBA). 4-HBA can serve as precursor for the manufacturing of nutraceuticals (e.g. coenzyme Q10, gastrodin, resveratrol), cosmetic ingredients (e.g. arbutin), drugs (e.g. xiamenmycin and shikonin), polymer fibers such as Vectran™, platform chemicals such as muconic acid, and deep eutectic solvents (Wang et al., 2018; Lin and Eudes, 2020). An approach for *in-planta* overproduction of 4-HBA consists in the expression of *Escherichia coli* chorismate pyruvate-lyase (UbiC) targeted to plastids to reroute chorismate away from the shikimate pathway. For example, sugarcane expressing *ubiC* contains up to 0.69% 4-HBA on a dry weight (DW) basis in leaves, which accumulates mostly as 4-HBA phenolic glucoside (McQualter et al., 2005).

In this work, we transformed the sorghum inbred line BTx430 with a construct containing *ubiC* as well as the gene encoding a feedback-resistant version of *E. coli* 3-deoxy-D-arabino-heptulonate-7-phosphate synthase (AroG*) in an attempt to increase the carbon flux through the shikimate pathway (**Figure 1a**). In this construct, both *E. coli* genes are preceded with the sequence of a plant signal peptide for targeting UbiC and AroG* to plastids (**Figure 1b**). The promoter of the rice polyubiquitin gene (*pRubi2*) was used to drive *ubiC* expression constitutively whereas the promoter of a maize cellulose synthase gene (*pZmCesa10*) involved in the formation of secondary cell walls was selected for *aroG\** expression to avoid possible toxicity and sterility issues (Oliva et al., 2021) (**Figure 1b**).

**Figure 1.**
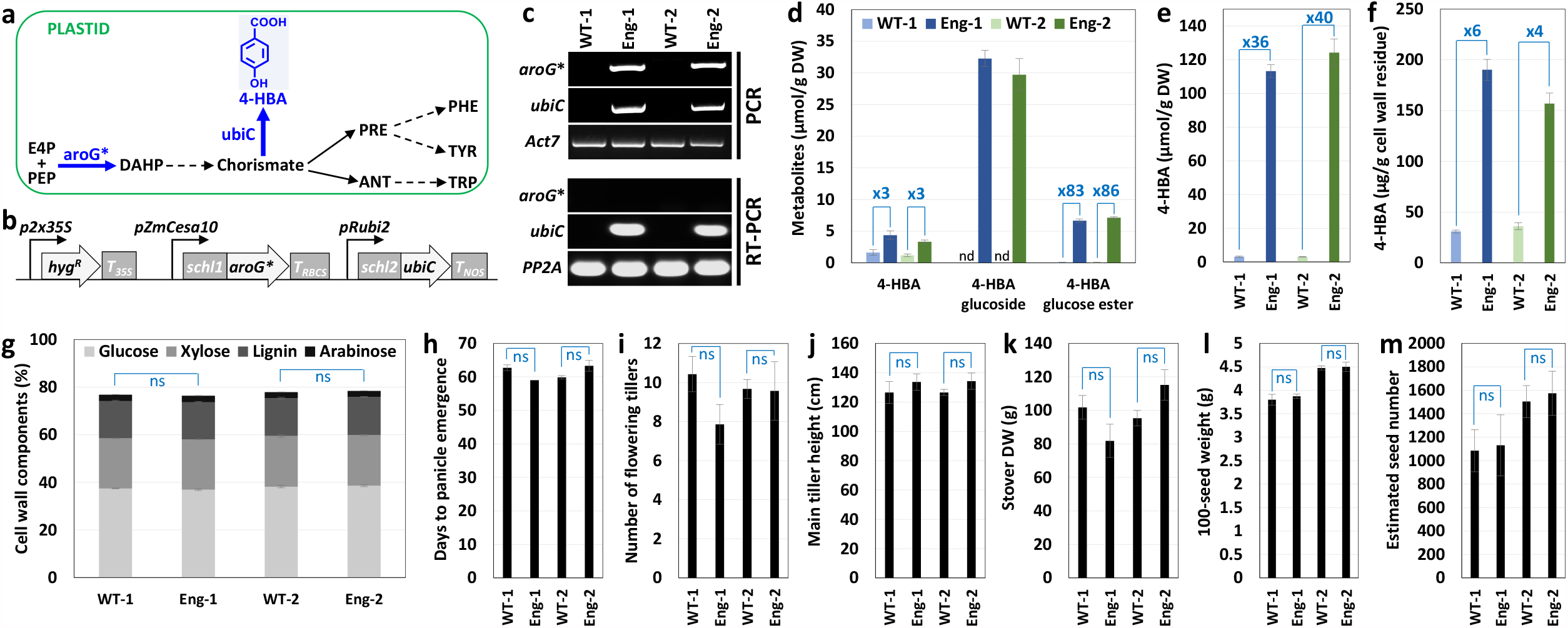
Overproduction of 4-HBA in sorghum. **(a)** Diagram of the shikimate pathway and engineering approach for 4-HBA overproduction. The two *E. coli* enzymes targeted to plastids are 3-deoxy-D-arabino-heptulosonate (DAHP) synthase with L175Q mutation (AroG*\**) and chorismate pyruvate-lyase (UbiC). Abbreviations: ANT, anthranilate; E4P, erythrose 4-phosphate; PEP, phosphoenolpyruvate; PHE, phenylalanine; PRE, prephenate; TRP, tryptophan; TYR, tyrosine. **(b)** DNA construct used for sorghum transformation. *Schl1* and *schl2* encode transit peptides from pea and maize ribulose-1,5-bisphosphate carboxylase (RuBisCo) small subunits, respectively. *pZmCesa10* and *pRubi2* designate the promoters of maize cellulose synthase 10 and rice polyubiquitin 2 genes. *p2×35S* is the enhanced 35S promoter from cauliflower mosaic virus (CaMV). *T*_*35S*_, *T*_*RBCS*_, and *T*_*NOS*_ are the terminators from CaMV 35S, Arabidopsis RuBisCo small subunit, and *Agrobacterium* nopaline synthase genes, respectively. *Hyg*^*R*^ denotes the aminoglycoside phosphotransferase marker gene used for plant selection. The jStack cloning method was used for gene assembly (Shih et al., 2016) and plasmid sequences are available at http://public-registry.jbei.org. The construct was transferred to sorghum via *Agrobacterium*-mediated transformation of calli obtained from immature embryos. **(c)** Molecular characterization of two independent engineered lines (Eng-1 and Eng-2). Upper panels: detection by PCR of *aroG*\* and *ubiC* using gDNA. Lower panels: detection of *ubiC* transcripts by RT-PCR using cDNA synthesized from mRNA obtained from stems of 4-week-old plants. *Act7* and *PP2A* sorghum genes were used to assess the quality of gDNA and cDNA, respectively. Wild-type segregants (WT-1 and WT-2) were used as negative controls. **(d)** Titers of 4-HBA and its glucose conjugates extracted from Eng-1 and Eng-2 using aqueous methanol. Twenty milligrams of biomass was sequentially extracted with 80% (v/v) methanol:water at 70°C and the resulting extracts were filtered and analyzed by LC-MS as previously described (Eudes et al., 2012; 2013). Metabolites were quantified by comparison with authentic standards. nd, not detected. **(e)** Total extractable 4-HBA content in lines Eng-1 and Eng-2. An aliquot of the methanolic extracts was dried in vacuo and treated with a 1 N HCl solution for 3h at 90°C. After partitioning with ethyl acetate, the amount of free 4-HBA recovered from the organic phase was quantified using LC-MS previously described (Eudes et al., 2012; 2013). **(f)** Cell-wall bound 4-HBA in lines Eng-1 and Eng-2. Extractive-free cell wall residues were treated with a 2 N NaOH solution for 24h at 30°C to release 4-HBA, followed by ethyl acetate partitioning and LC-MS analysis as previously described (Eudes et al., 2012; 2013). **(g)** Cell wall composition of lines Eng-1 and Eng-2. Monosaccharides and lignin were measured using the Klason method as previously described (Eudes et al., 2012). (**h-m**) Agronomic parameters of lines Eng-1 and Eng-2 compared to wildtype controls. Fully mature senesced plants were used for i-m.

Two independent lines (Eng-1 and Eng-2) containing the dual *aroG\**-*ubiC* construct and their respective wild-type segregants (WT-1 and WT-2) were selected for characterization in the T2 generation. Although we detected by PCR the presence of *aroG\** and *ubiC* in both Eng-1 and Eng-2, only the expression of *ubiC* could be confirmed in these lines using RT-PCR (**Figure 1c**). Whether the absence of *aroG*\* expression is due to a gene silencing effect or to the lack of *pZmCesa10* activity in sorghum remains uncertain. Next, fully mature senesced dry plants were harvested without their panicles and ground to a fine powder for metabolite analyses. Using liquid chromatography-mass spectrometry (LC-MS), measurements of free 4-HBA and potential 4-HBA glucose conjugates were analyzed in methanolic extracts obtained from biomass samples. A small amount of free 4-HBA was detected in wild-type extracts (1.2–1.6 µmol/g DW) and increased 3-fold in engineered lines (**Figure 1d**). 4-HBA phenolic glucoside, which was undetectable in wild-type extracts, reached 32.3 and 29.7 µmol/g DW in extracts from lines Eng-1 and Eng-2, respectively (**Figure 1d**). Furthermore, 4-HBA glucose ester was detected and increased 83-and 86-fold in transgenics, which corresponds to titers of 6.6 and 7.1 µmol/g DW (**Figure 1d**). Finally, an aliquot of methanolic extracts was subjected to acid hydrolysis in order to release 4-HBA from its conjugated forms. Measurement of 4-HBA aglycone in resulting hydrolysates showed that transgenic lines Eng-1 and Eng-2 contain 113 and 124 µmol 4-HBA/g DW, respectively (**Figure 1e**). These values are equivalent to 4-HBA titers of 1.56% and 1.72% DW, which represents a 36-and a 40-fold increase, respectively, compared to the titer measured in wildtypes (**Figure 1d**). Since the total amount of free 4-HBA and 4-HBA glucose conjugates in methanolic extracts is below that of 4-HBA aglycone measured in acid-treated extracts, we conclude that other unidentified 4-HBA conjugate forms accumulate in transgenics. Finally, free 4-HBA was measured in alkaline hydrolysates obtained from extractive-free cell wall preparations. Compared to controls, six and four times more 4-HBA was released from Eng-1 and Eng-2 cell walls, which represents 190 and 157 µg 4-HBA/g cell wall residue, respectively (**Figure 1f**). Although 4-HBA ester end-groups have been described in lignin fractions from willow, poplar, aspen, Neptune grass, and palms, the exact mode of 4-HBA attachment to cell walls in sorghum remains to be determined.

The shikimate pathway provides aromatic amino acids (i.e. phenylalanine, tyrosine, and tryptophan) important for several physiological functions during plant development (**Figure 1a**). Therefore, we carried out several measurements of biomass composition and agronomic traits to evaluate possible growth defects in engineered lines. Cell walls, which are the main constituents of plant biomass, showed no significant difference between transgenics and controls regarding the proportion of major components including glucose (representing cellulose and mixed-linkage glucan), xylose and arabinose (representing xylan), and lignin (**Figure 1g**). Moreover, no differences were observed for several developmental and yield parameters including the number of days to panicle emergence (**Figure 1h**), number of flowering tillers and height of the main tiller (**Figures 1i** and **1j**), stover biomass yields (**Figure 1k**), and seed weight and number of seeds per plant (**Figures 1l** and **1m**). Overall, engineered sorghum overproducing 4-HBA shows no penalty yield nor developmental defects while grown under controlled environment in the greenhouse.

To conclude, this work demonstrates the transferability of the *ubiC* engineering approach to sorghum and obtained engineered lines accumulate up to 1.7% DW of the valuable bioproduct 4-HBA. Considering that several important metabolites derive from chorismate and more generally from the shikimate pathway, it will be important to field test these lines to assess their agronomic performance and resilience to environmental stress under natural conditions.

Values are means ±SE of six biological replicates and significant differences observed between transgenics and wildtypes have *P* values less than 0.001 using the unpaired Student’s t-test. ns: no significant difference.

## Acknowledgements

This work conducted by the Joint BioEnergy Institute was supported by the US Department of Energy, Office of Science, Office of Biological and Environmental Research under contract no. DE-AC02-05CH11231 between Lawrence Berkeley National Laboratory and the US Department of Energy. Part of the work was supported by the Laboratory Directed Research and Development Program of Lawrence Berkeley National Laboratory (J.T. and T.R.N). Authors are thankful to Professor Kazufumi Yazaki (Kyoto University, Japan) for providing the 4-hydroxybenzoate glucose conjugate standards and to Dr. Roger Thilmony (USDA, Western Regional Research Center, Albany, CA) for providing the *pRubi2* promoter.

## Conflict of interest

The authors have no conflict of interest to declare

## Author contributions

C-YL and JT performed DNA clonings. APK, JH, and KN-V transformed sorghum. MYL and YT genotyped the plants. C-YL conducted gene expression analyses. YT grew the plants and measure agronomic traits. YT did the metabolite and cell wall analyses. RK and EEKB performed LC-MS analyses. AE wrote the paper. APK, TRN, EEKB, HVS, and AE supervised the research. All authors read and approved the final manuscript.

## Notes

### Competing Interest Statement

The authors have declared no competing interest.

